# A framework for integrating genomic profiles into captive breeding and reinforcement programmes: A case study on captive saker falcons

**DOI:** 10.1101/2025.07.04.663124

**Authors:** TB Hoareau, A Barbosa, X Velkeneers, G Leveque, L Lesobre

**Affiliations:** Reneco International Wildlife Consultant LLC, Abu Dhabi, United Arab Emirates

**Keywords:** Conservation breeding, Genetic Diversity, Inbreeding, *Falco cherrug*, Wild reinforcement, Whole genome sequencing

## Abstract

Ex-situ conservation is crucial for preserving endangered species by breeding to preserve genetic diversity and provide surplus for translocation, thereby supporting in-situ conservation and enhancing wild populations. Genomic tools can assist breeding strategies and ensure long-term success of ex-situ conservation efforts by assessing genetic introgression, determining genetic origin and status, and inferring genetic relatedness of potential founders. This study aims to develop a comprehensive genomic approach for assessing the genetic profiles of candidate founders for ex-situ breeding, with the goal of releasing surplus individuals while using the endangered Saker Falcons (*Falco cherrug*) as a study model. Genetic clustering of 31 captive sakers revealed both diverse origins, some matching wild Asian individuals (Mongolia), and a lineage (Group III) divergent from wild populations. Comparative analyses detected hybridisation signals in 61.3% of individuals, including three with gyrfalcon (*F. rusticolus*) introgression and Group III’s pronounced divergence indicating past interbreeding with an unknown falcon species. All captive birds exhibited severe inbreeding (FROH = 0.352), far exceeding wild population levels (FROH = 0.131). Using the partial pedigree data of the captive sakers, we established a genetic relatedness threshold of 0.154 (95% CI: 0.096–0.211) to identify cases of related dyads (both full and half-siblings). At this threshold, 18.3% of captive dyads showed relatedness, with asymmetric genetic contributions between pairs, reflecting a functionally small breeding flock. To avoid risks from releasing admixed or inbred individuals, we recommend excluding introgressed birds, strategically pairing purebreds, and sourcing new founders from genetically validated wild sources, especially underrepresented Central Asian lineages. Applying this genomic framework, we demonstrate its role in safeguarding genetic integrity and preventing genetic erosion in conservation breeding programmes, thereby establishing a standard for genomic-led ex-situ conservation.

## Introduction

The conservation of threatened species often requires a multifaceted approach that integrates both in-situ and ex-situ management strategies (Gant et al., 2021; Maxted, 2013; Schwartz et al., 2017). A cornerstone of this integration is leveraging captive-bred individuals for conservation translocations to demographically support in-situ populations. However, the success of such programmes depends on the genetic characteristics and origin of captive-bred individuals (Frankham, 2008; Jamieson & Lacy, 2012; Tracy et al., 2011). Their genomic profiles play a pivotal role in determining their suitability for release, as genetic mismatches can lead to significant ecological and evolutionary consequences (Frankham, 2008; Ortego et al., 2024). For instance, if founders are genetically too divergent from the recipient populations, whether due to adaptively different lineages or hybridisation with other taxa, the risk of outbreeding depression increases (Frankham et al., 2010). This can result in maladaptation, loss of critical adaptive traits, and disruption of co-adapted gene complexes (Edmands, 2007; Frankham, 2015). Conversely, if the individuals are closely related, inbreeding depression may ensue, leading to reduced genetic diversity, increased expression of deleterious alleles, and a decline in overall fitness of the progeny (Frankham et al., 2019). Both scenarios, outbreeding and inbreeding depression, pose serious threats to the viability of wild populations, with potentially unpredictable and detrimental outcomes (Frankham et al., 2010). Therefore, ex situ populations require rigorous genomic evaluation to ensure their genetic integrity aligns with conservation goals, avoiding the issues of outbreeding and inbreeding depression. Therefore, establishing a dedicated framework to genomically evaluate ex situ populations is essential to assess their genetic integrity, origin, and status, thus ensuring the long-term success and sustainability of translocation programmes.

To address this and mitigate potential genetic risks, we propose a comprehensive genomic framework that: (1) screens for hybridization, (2) assigns population origins, and (3) quantifies genetic health—criteria essential for selecting optimal founder candidates (Ballou & Lacy, 1995). First, this framework should screen for hybridisation and introgression using diagnostic markers (e.g., SNPs) to exclude individuals with admixed ancestry that could disrupt local adaptation (Fitzpatrick et al., 2020; VonHoldt et al., 2022). Second, it must determine the population of origin of captive individuals through assignment tests (e.g., PCA, STRUCTURE, or DAPC) to ensure alignment with the genetic background of target wild populations (Manel et al., 2003; Puckett & Eggert, 2016). Finally, the framework should assess genetic status by quantifying individual genome-wide diversity (e.g., heterozygosity), inbreeding coefficients (e.g., FROH), and relatedness among potential breeders (Kardos et al., 2016). By integrating these three pillars, hybrid detection, origin assignment, and genetic health assessment, such a framework empowers conservationists to identify individuals that maximise adaptive potential while minimising risks of outbreeding or inbreeding depression, thereby safeguarding the ecological and evolutionary integrity of reinforced populations (Kardos et al., 2021).

The saker falcon (*Falco cherrug*), a globally threatened apex predator of Eurasian grasslands and steppes, represents the complex interplay between ex-situ conservation and in-situ recovery. Its decline disrupts trophic cascades (Ferguson-Lees & Christie, 2001), while its unique adaptations, including thermal resilience in arid environment, and high-altitude specialisation, make it evolutionarily unique (Pan et al., 2017; Zhan et al., 2013). This cultural keystone species has been used in falconry traditions for millennia (Stretesky et al., 2018), further underscoring its anthropogenic significance. Catastrophic declines (60% in key populations) linked to habitat loss and harvesting (Karyakin et al., 2023) have led to functional extinctions in some regions (Ragyov et al., 2014), while European groups continue to struggle with the genetic legacy of DDT crashes (BirdLife International, 2017).

While captive breeding offers crucial demographic support for threatened saker populations, current programmes face significant genetic management challenges, particularly due to the absence of standardised protocols addressing three critical issues: (1) undetected introgression with closely related species, a persistent problem in captive flocks, especially with gyrfalcons that mtDNA alone cannot resolve (Nittinger et al., 2007); (2) mismatched population origins, which may introduce maladaptive traits (Besnier et al., 2022); and (3) unchecked inbreeding and eroded diversity, that could compromise long-term viability (Kardos et al., 2021). These gaps are exacerbated by missing wild reference data, as seen in Eastern European captive populations ( e.g., Bulgaria; Petrov et al., 2023), underscoring the urgency of developing genomic tools for informed decision-making in saker ex-situ conservation.

Recent genomic advances have overcome previous limitations. While microsatellites could distinguish sakers from gyrfalcons (Johnson et al., 2007), whole-genome sequencing (Hu et al., 2022) now enables far greater precision in hybrid detection, population assignment and evaluation of individual genetic diversity and inbreeding levels (Zuccolo et al., 2023). Here, we apply the proposed framework to captive sakers, aiming to: (1) diagnose gyrfalcon introgression in the captive flock, (2) conduct population assignment to differentiated wild origins (Eastern Europe, Mongolia, Tibetan Plateau), and (3) assess their suitability as breeders to produce individuals for the reinforcement of wild populations in Kazakhstan. Our approach addresses urgent saker conservation needs while providing a replicable blueprint for genomic screening in other conservation breeding programmes (Badia-Boher et al., 2022).

## Material and methods

### Samples, genomic library and re-sequencing

A total of 31 saker falcons, sourced from private captive breeding facilities, are currently maintained in the Sheikh Khalifa Houbara Breeding Centre in Kazakhstan (i.e. SKHBC-KZ; Table 1). Blood samples were collected and preserved in 95% ethanol. Blood sampling procedures followed national regulations and institutional ethical guidelines, minimising stress and harm to the birds. Necessary permits were obtained, ensuring full compliance with legal and ethical standards and birds were housed under conditions meeting animal welfare standards.

**Table 1.** Sample details of the 31 evaluated saker falcons. Line colour shadings indicate individuals sharing the same parents, with different coloured cells representing the parents of these individuals.

| INDIVIDUAL ID | SEX | MOTHER | FATHER | PARENT PAIR<br>ID |
| --- | --- | --- | --- | --- |
| F11K00001 | M | Unk | Unk | NA |
| F12K00001 | F | Wild | Wild | NA |
| F13K00002 | M | Wild | Wild | NA |
| F14K00001 | F | Unk | Unk | NA |
| F14K00002 | F | Wild | Wild | NA |
| F15K00002 | M | Wild | Wild | NA |
| F16K00001 | M | Unk | Unk | NA |
| F16K00002 | M | Unk | Unk | NA |
| F16K00003 | M | Unk | Unk | NA |
| F17K00003 | M | Unk | Unk | NA |
| F20K00002 | F | Unk | Unk | NA |
| F20K00003 | F | Unk | Unk | NA |
| F20K00004 | M | Unk | Unk | NA |
| F20K00005 | F | Unk | Unk | NA |
| F20K00006 | F | Unk | Unk | NA |
| F20K00007 | M | Unk | Unk | NA |
| F20K00008 | F | Wild | Wild | NA |
| F11M00074 | F | F04M00004 | F03M00015 | 1 |
| F12M00109 | M | F04M00004 | F03M00015 | 1 |
| F13M00018 | M | F04M00004 | F03M00015 | 1 |
| F13M00098 | F | F04M00004 | F03M00015 | 1 |
| F13M00140 | M | F05M00027 | F03M00015 | 2 |
| F13M00054 | M | F05M00027 | F06M00023 | 3 |
| F14M00063 | M | F05M00027 | F07M00022 | 4 |
| F12M00107 | F | F06M00025 | F99M00057 | 5 |
| F13M00103 | M | F06M00025 | F99M00057 | 5 |
| F14M00065 | M | F06M00025 | F99M00057 | 5 |
| F13M00082 | F | F09M00082 | F09M00004 | 6 |
| F14M00022 | F | F09M00082 | F09M00004 | 6 |
| F14M00077 | F | F09M00082 | F09M00004 | 6 |
| F13M00149 | F | F10M00119 | F03M00015 | 7 |

Genomic DNA was extracted either by phenol chloroform or using spin columns following the NEB Monarch Genomic DNA Purification Kit Protocol (New England Biolabs, 2022). Short read sequence data was generated on Illumina NovaSeq at the Oklahoma Medical Research Foundation (OMRF) yielding reads with a length of 150 bp. All samples were sequenced with an expected range of 20–30× coverage.

### Genomic data

We used the chromosome-level gyrfalcon genome assembly (Zuccolo et al., 2023; GCA_028534605.1) as the reference due to its superior quality (33 haploid chromosomes) and close evolutionary relationship with saker falcons (Johnson et al., 2007). We compared genomic data from captive individuals to wild reference samples representing three genetically distinct saker populations that were previously studied (Eastern Europe, Mongolia, Tibetan Plateau; Hu et al., 2022; Figure 1).

**Figure 1.**
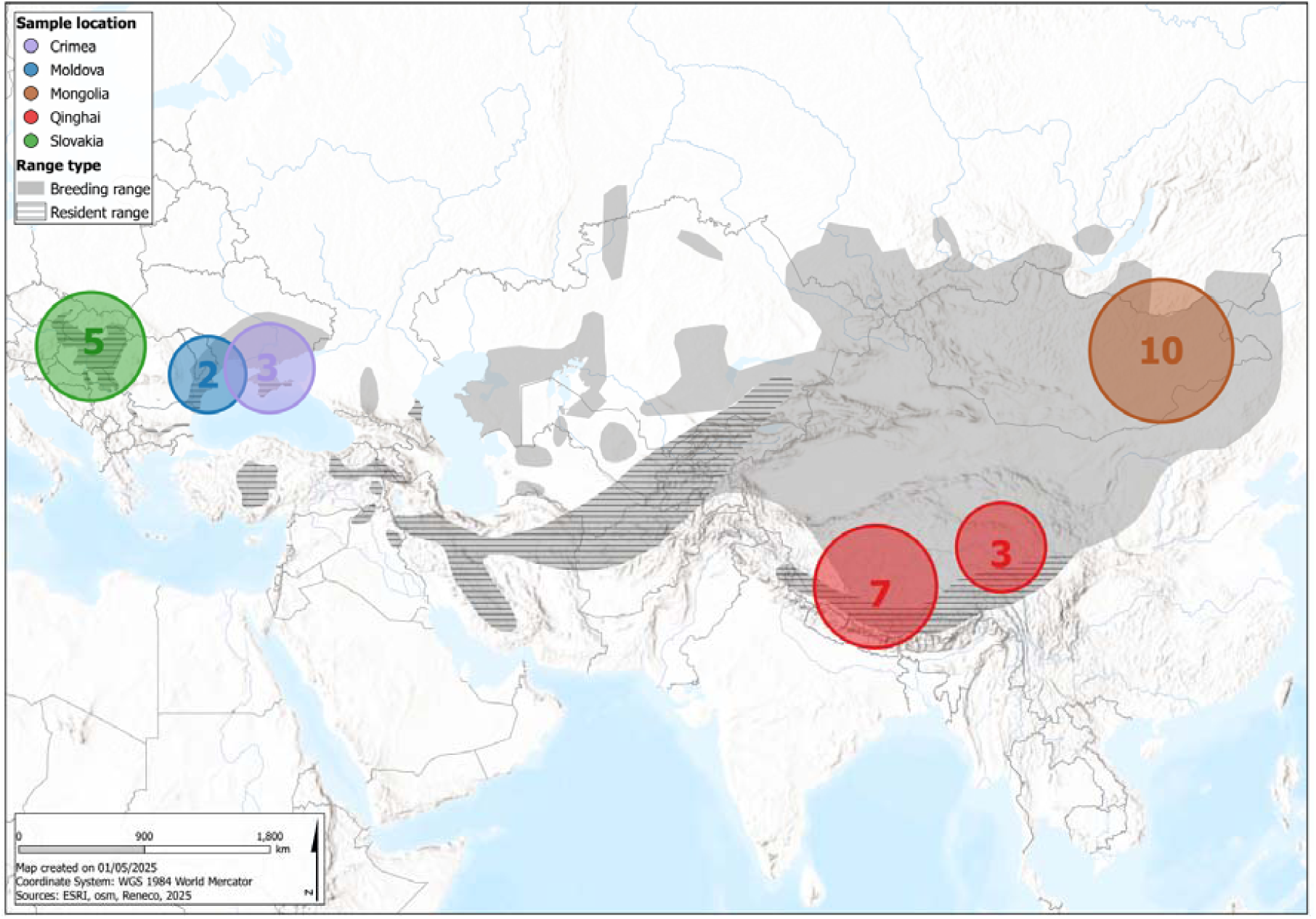
Distribution range of Saker falcons, with sampling locations (circles = sample size). Central Asia (highlighted) represents a critical but under-sampled region with high habitat fragmentation.

### Data processing

Reads were quality-checked and duplicates removed using FASTP (v0.20.0; https://github.com/opengene/fastp) then mapped to the gyrfalcon genome with BWA-MEM (Li & Durbin, 2009). The resulting BAM files were processed using SAMTOOLS (Li et al., 2009). Variant calling was then performed using SAMTOOLS mpileup and BCFTOOLS (v1.17) and variants were filtered using BCFTOOLS and VCFTOOLS (Danecek et al., 2011) using the following criteria: mapping and genotype phred scores ≥20, biallelic SNPs only, and sequencing depth within the 5^th^-95^th^ percentile range.

#### Data analyses

PCA was performed using PLINK v1.90 (Purcell et al., 2007) following linkage disequilibrium pruning. First, we included all individuals (including gyrfalcons) to assess hybridisation. Second, we analysed saker falcons against reference populations to determine their origins. Using Mclust (Fraley & Raftery, 2002) on PCA coordinates, we leveraged multidimensional genetic variance to identify the number of genetic cluster and verify the assignment of captive sakers to reference populations. This model-based algorithm evaluates and compares 14 covariance models (EEE-VVV) across k=1–9 clusters, comparing cluster form and structures while calculating assignment probabilities.

We calculated observed (*Ho*) and expected (*He*) heterozygosity using BCFTOOLS v1.17 to assess genetic diversity in captive and wild saker falcons, with inbreeding coefficients (F) derived from the same analysis. In addition, we identified ROH (>100 kb) using PLINK v1.90 with the following parameters: a minimum of 50 consecutive homozygous SNPs, a SNP density ≥1 SNP per 100 kb, and a 50-SNP sliding window allowing up to 3 heterozygous or 5 missing SNPs per window, shifting one SNP at a time. FROH were obtained by dividing the total ROH length by the reference genome assembly size (1.2 TB). Wild individuals provided baseline ROH values for comparison with captive birds.

Pairwise relatedness was estimated using the RAB coefficient (NGSRELATE V2; Hanghøj et al., 2019), which calculates identity by descent proportions while accommodating inbreeding. Unlike standard kinship coefficients, RAB remains robust under population stratification, making it particularly suitable for captive populations with possible mixed origins and varying inbreeding levels (Hedrick & Lacy, 2015). Using known parent-offspring pairs from captive pedigrees, we established a relatedness threshold by: (1) calculating values for sibships (≥1 shared parent) and unrelated wild individuals (different locations), then (2) fitting a logistic regression model (R package aod v1.3) to distinguish these groups. This threshold is used to inform pairing strategies to avoid pairing related birds.

## Results

### Family structure of the breeders

In our dataset of 31 individuals, only 14 have pedigree records available (Table 1). These records have allowed us to establish seven distinct families based on shared parent pairs, with some parents contributing to multiple individuals. For instance, male F03M00015 is the father of six current breeders, while males F99M00057 and F09M00004 have each fathered three of the breeders. Similarly, several females have contributed to multiple individuals, with individual F04M00004 mothering four of the breeders, and individuals F05M00027, F06M00025 and F09M00082 each mothering three breeders. Taking this into account, our dataset comprises 12 full-sib and 13 half-sib combinations, but no parent-offspring. Apart from these cases, the relationship levels among dyads of the captive sakers remains poorly characterised due to limited pedigree data.

### Data generation and processing

Whole-genome sequencing of the 31 captive sakers yielded 529 GB of data (mean 17.1 GB/individual), with 247 million clean reads per individual (range: 190-340 million). The dataset showed moderate duplication (mean 9.1%, range 3.4 – 12.6%) and 24× coverage on average (Table S2). The final dataset contained 4 538 584 autosomal polymorphic SNPs in the captive birds. A total of 116 466 SNPs remained after linkage disequilibrium pruning for the PCA analyses.

### Genetic structure and assignment of captive sakers

PCA demonstrates clear genetic differentiation between wild sakers and gyrfalcons along PC1, which explains the greatest proportion of variance in the dataset (Figure 2A). Three captive individuals (F14K00001, F16K00001, F17K00003) exhibit intermediate clustering between Mongolian sakers and gyrfalcons, consistent with gyrfalcon introgression, while the remaining individuals display substructure.

**Figure 2.**
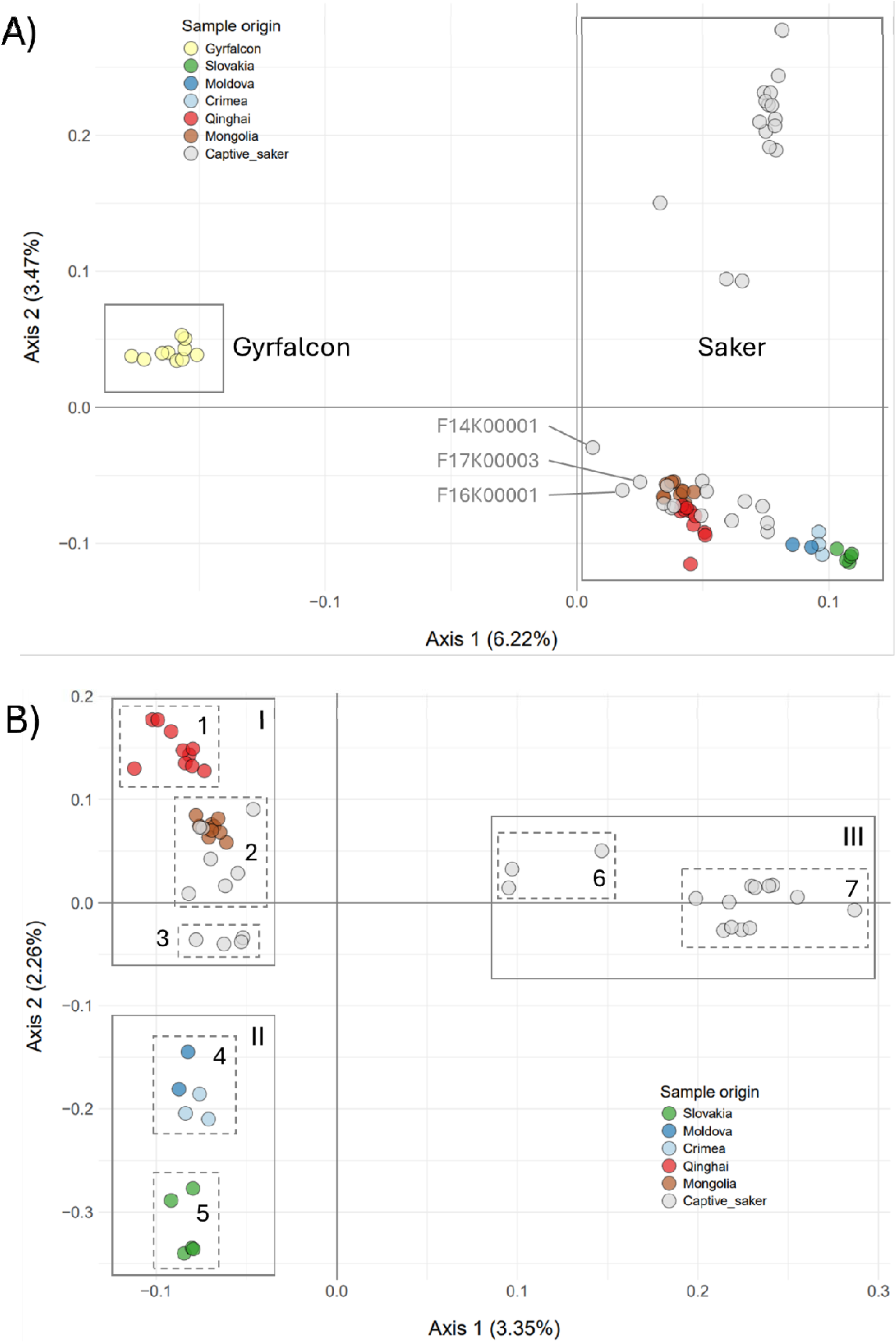
Genetic structure of captive and wild saker falcons revealed by Principal Component Analysis. (A) Captive individuals (light grey) compared to wild saker (other colours) and gyrfalcon (yellow) reference populations. (B) Captive individuals compared to wild saker reference populations representing three genetic groups and seven clusters: Group I (Central Asia/Captive sakers), Group II (Eastern Europe), and Group III (Captive sakers of unknown origin).

Joint analysis of PCA and clustering results identifies significant hierarchical population structure in Saker falcons. At a larger scale, three main clusters are identified (Figure 2B; Figure S1): Group I includes wild Saker Falcons from Mongolia and Qinghai and some captive birds, including the three birds presenting signs of introgression with gyrfalcons (N = 15); Group II comprises wild Saker Falcons from Eastern Europe only, without any captive sakers; and Group III (N = 16) consists exclusively of captive Saker Falcons that cluster separately from wild Sakers. These groups further subdivide into seven subgroups. Group I splits into Subgroup 1, which includes wild Saker Falcons from the Qinghai-Tibet Plateau, and Subgroup 2, consisting of Mongolian Saker Falcons and 10 SKHBC captive birds. Subgroup 3 comprises captive Saker Falcons from SKHBC only (N = 5). Group II divides into Subgroup 4, which includes birds from Crimea and Moldova (grouped as one cluster hereafter), and Subgroup 5, which includes birds from Slovakia. Group III comprises Subgroup 6, which includes three captive birds, and Subgroup 7, which includes 13 captive birds.

#### Genetic diversity and inbreeding levels

Captive sakers exhibit lower observed heterozygosity (mean HOBS = 0.126 ± 0.008) compared to wild populations (mean HOBS = 0.143 ± 0.022), though with partial overlap in their value ranges (Table 2; Figure 3). The analysis of Runs of Homozygosity (ROH) revealed contrasting patterns between captive and wild Saker Falcons (Table 2). Notably, wild individuals had an average FROH of 0.131 ± 0.060, some exhibiting low (< 0.1) and others intermediate (0.1–0.2) proportion of genome in ROH (i.e. ROH-derived inbreeding coefficient FROH), indicating a relatively consistent level of inbreeding within these wild populations. In stark contrast, captive sakers exhibited significantly higher genomic inbreeding levels (FROH = 0.352 ± 0.024; Figure 3) that showed no overlap with wild population values. These patterns reflect widespread homozygosity in the captive birds, confirming both reduced genetic diversity and elevated inbreeding. Most notably, individual F14K00001 showed extreme genomic signatures - the lowest heterozygosity (HOBS = 0.100) and highest inbreeding level (FROH = 0.420) of all captive birds, indicating particularly severe genetic erosion.

**Figure 3.**
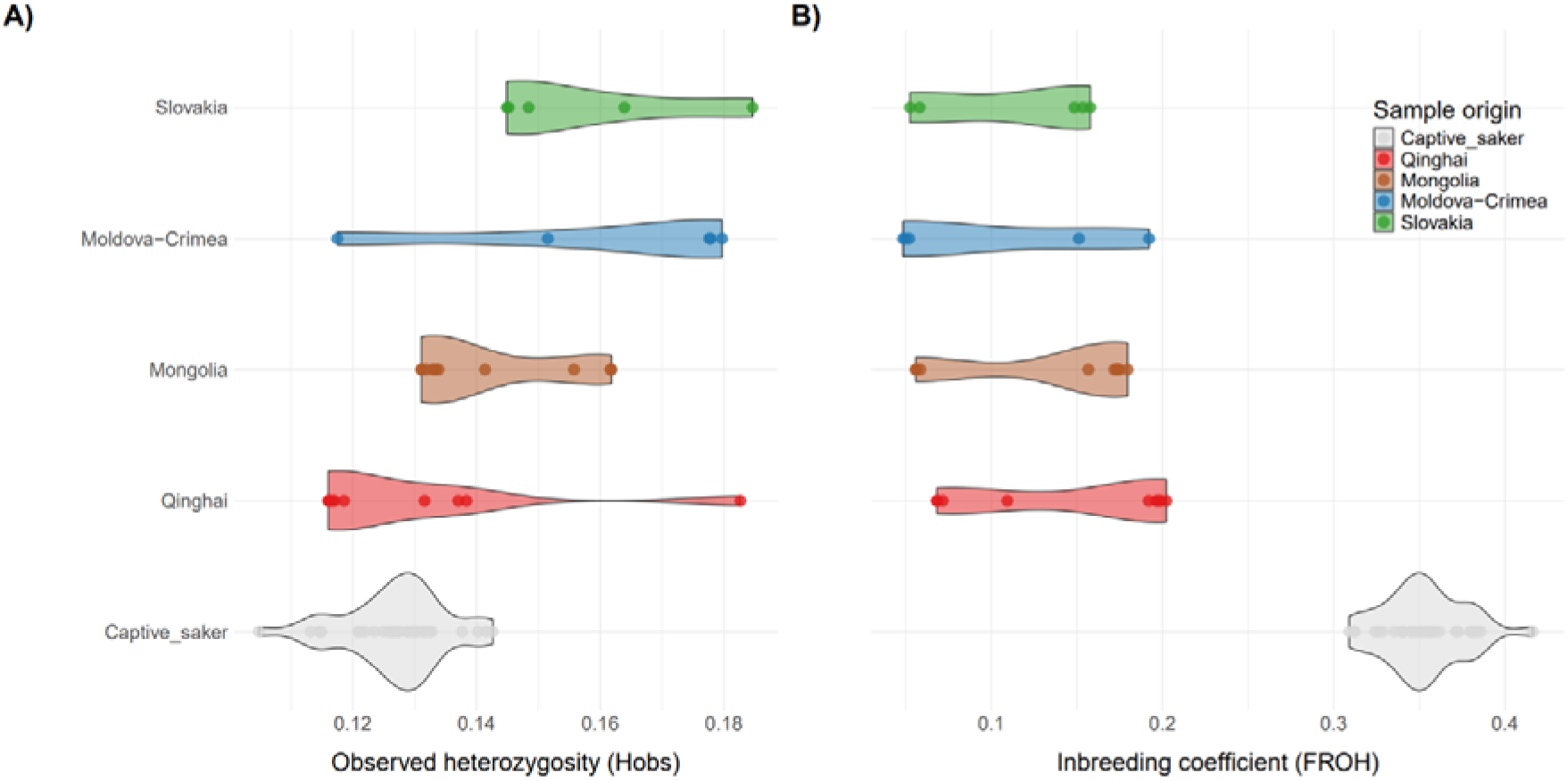
Comparison of (A) genetic diversity and (B) inbreeding levels (i.e. proportion of the genome in runs of homozygosity or ROH) between captive sakers and wild individuals from reference populations. Captive sakers generally show lower genetic diversity and higher ROH values (0.3-0.4 of their genome) compared to wild individuals (0-0.2 of their genome). Notably, individual F14K00001 has the highest ROH value at 41.6%.

**Table 2.** Genetic diversity, inbreeding levels and relatedness values for wild and captive saker falcons across different geographic regions.

| Sample category | Region | *N* | Observed heterozygosity (*H<sub>obs</sub>*) | Inbreeding level (FROH) | Relatedness (*R<sub>AB</sub>*) |
| --- | --- | --- | --- | --- | --- |
| *Wild individuals* |  |  |  |  |  |
| Slovakia | Eastern Europe | 5 | 0.157 ± 0.017 | 0.114 ± 0.053 | 0.250 ± 0.084 |
| Crimea-Moldova | Eastern Europe | 5 | 0.160 ± 0.026 | 0.098 ± 0.068 | 0.147 ± 0.045 |
| Mongolia | Central-eastern Asia | 10 | 0.141 ± 0.013 | 0.137 ± 0.055 | 0.058 ± 0.016 |
| Qinghai | Central-eastern Asia | 10 | 0.129 ± 0.020 | 0.150 ± 0.062 | 0.142 ± 0.012 |
| Overall wild |  | 30 | 0.143 ± 0.022 | 0.131 ± 0.06 | 0.054 ± 0.057 |
| *Captive individuals* |  |  |  |  |  |
| Group I |  | 15 | 0.129 ± 0.009 | 0.345 ± 0.028 | 0.029 ± 0.093 |
| Group II |  | 0 | – | – | – |
| Group III |  | 16 | 0.125 ± 0.007 | 0.360 ± 0.018 | 0.032 ± 0.080 |
| Overall captive |  | 31 | 0.126 ± 0.008 | 0.352 ± 0.024 | 0.069 ± 0.136 |

#### Relatedness and evaluation of the pairing strategy

Captive sakers showed higher and more variable relatedness values (mean *RAB* = 0.069 ± 0.136; range: 0.000 – 0.696) compared to wild conspecifics (mean *RAB* = 0.054 ± 0.084; range: 0.000–0.394; Figure 4A), confirming the presence of closely related individuals in this captive flock. Wild individuals presented regional variation, with Mongolia displaying the lowest relatedness (mean *RAB* = 0.058 ± 0.016; range: 0.025–0.094), followed by Qinghai-Tibet (mean *RAB* = 0.142 ± 0.012; range: 0.110–0.187) and Eastern European birds from Crimea/Moldova (mean *RAB* = 0.147 ± 0.045; range: 0.083–0.234). Notably, wild Slovakian individuals showed elevated relatedness (mean *RAB* = 0.250 ± 0.084; range: 0.083–0.234).

**Figure 4.**
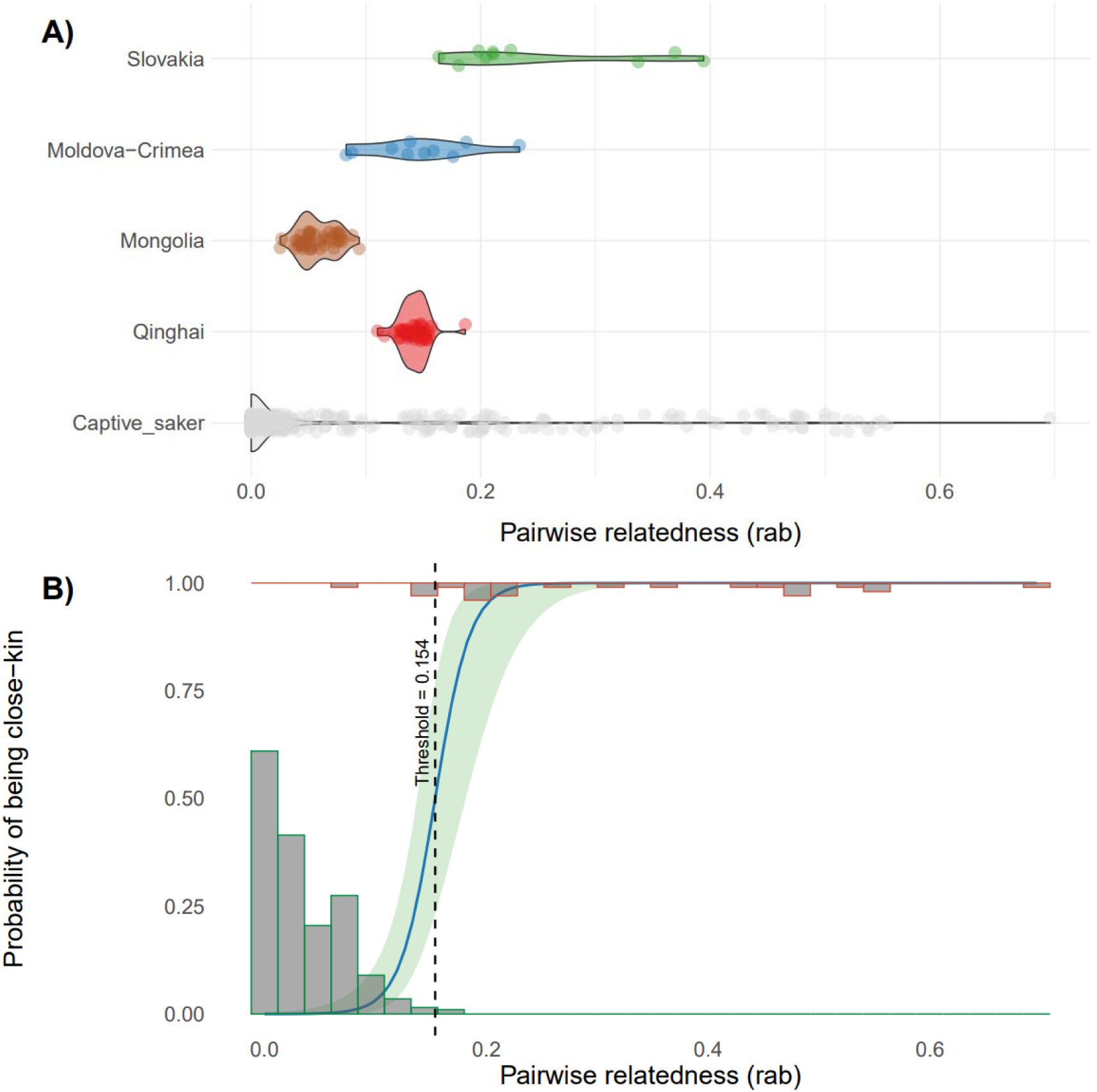
Genetic relatedness patterns in captive and wild Saker falcons. (A) Within-population relatedness (RAB) distributions, showing elevated values in captive birds (light grey) compared to wild reference populations (coloured by geographic origin). (B) Logistic regression of kinship probability against relatedness values (Wald test, χ*²* = 18.4, *df* = 1, *P* = 1.8 × 10⁻D), with histograms showing the distribution of unrelated (green) and related (red) dyads. The dashed line indicates the RAB threshold (0.154) that best discriminates between these groups.

To develop a tool for identifying related individuals and informing optimal pairing strategies, we assessed relatedness between known parent-offspring dyads (mean *RAB* = 0.321 ± 0.171; including full-sibs and half-sibs from captive pedigrees) and unrelated wild dyads (mean *RAB* = 0.034 ± 0.035; Figure 4B). The fitted logistic regression model significantly distinguishes these groups (Wald test: χ² = 18.4, *df* = 1, *P* = 1.8 × 10 D), yielding a relatedness threshold of 0.154 (95% CI: 0.096–0.211) above which individuals can be considered as related. This threshold identified 13.8% of related dyads among captive individuals with unknown pedigree (n = 61 out of 441 pairs), involving 20 individuals that contributed unevenly to these dyads (data not shown). Combined with captive individuals with known pedigree, this percentage increased to 18.3% of the dyads (n = 85 out of 465).

## Discussion

Applying our framework to the genomic assessment of captive Saker Falcons reveals two critical conservation challenges: (1) multiple divergent lineages within the captive population possibly resulting from widespread introgression from non-native species (affecting ∼52% of potential breeders), and (2) severe inbreeding (FROH > 0.35). This underscores the risks of releasing admixed or genetically impoverished individuals into wild populations, which could undermine local adaptations and exacerbate biodiversity loss. By integrating high-resolution SNP data with clustering and relatedness analyses, we demonstrate how genomic tools can diagnose these issues, providing a framework for evidence-based management of genetic diversity. Below, we explore 1) the origins and genetic integrity of the birds, including evidence of introgression, 2) key differences in genetic status (diversity, inbreeding and relatedness) between captive and wild individuals, and 3) actionable measures to develop conservation breeding to safeguard the species’ genetic integrity.

### 1. Genetic integrity and potential origin of captive sakers.

Maintaining genetic integrity in captive Saker Falcons is critical for the release programme, yet our study reveals signs of introgression. Using a combination of DAPC and MCLUST clustering, we identified low-level gyrfalcon ancestry in ∼9.7% of captive sakers (linked to Group I). Additionally, individuals from the Group III that represent 51.6% of the potential breeders may also present introgression from another unidentified source species (see discussion below). These findings reflect established falconry practices, where intentional hybridisation, particularly saker × peregrine and saker × gyrfalcon crosses, is routinely used to produce birds with desirable traits (Andrew Dixon, 2012; Eastham & Nicholls, 2005). Moreover, hybrids represent over a quarter (26.8%) of the global trade, although the purebred (gyrfalcons at 16.2% and sakers at 11.5%) retain higher market value (Panter et al., 2023).

While traditional mtDNA markers lack the resolution to detect hybridisation in hierofalcons due to incomplete lineage sorting (Nittinger et al., 2007), recent genomic analyses have confirmed clear autosomal differentiation between Saker and Gyrfalcons, with no signs of wild introgression for over 16,500 years (Hu et al., 2022). Mirroring these genomic findings, our study provides species-level differentiation while achieving the resolution necessary to distinguish pure from introgressed individuals. This precision tool is critical for conservation, as failure to identify admixed individuals, whether released accidentally or intentionally, may result in outbreeding depression, genetic swamping of wild populations, and the erosion of locally adaptive traits (Richard Frankham et al., 2010; Kovach et al., 2015). However, achieving this level of discrimination demands comprehensive reference data to reliably identify introgression and assess their conservation implications.

The exclusive presence of captive birds in Cluster III suggests two explanations, both contingent on available reference data: (1) origin from an unsampled saker population absent from current wild genetic references, or (2) introgression from another falcon species (excluding gyrfalcons) that remains undetected due to limited comparative samples. The first hypothesis aligns with documented saker subspecies diversity in Central Asia (Karyakin, 2011; Figure 1), a region notably underrepresented in genomic studies (Hu et al., 2022; Pan et al., 2017), potentially leaving Central Asian lineages unaccounted for. The population in Turkey (Dixon et al., 2009) might represent another such a cryptic group of divergent individuals. However, Cluster III (Figure 2B) shows striking genetic divergence, even more distinct from wild sakers than geographically remote populations like Mongolia and Qinghai birds, which remains closer to East Europeans sakers. This strongly points to human-mediated hybridisation with other falcons (e.g., peregrine or lanner falcons). Regardless of its origin, the absence of Cluster III in wild datasets raises critical questions about its evolutionary history and conservation risks, especially for conservation breeding and reintroduction programmes.

Besides the possible cases of introgression, some captive birds (N=10) cluster closely with wild Mongolian Sakers (Group I, Subgroup 2; Figure 2), suggesting they may represent authentic Asian lineages. This result is especially important as it highlights individuals with genetic profiles aligned to wild populations, making them candidates for potential conservation breeding. The intermediate clustering of five captive birds (Group I, Subgroup 3) between Mongolian and Eastern European populations may represent previously undescribed Central Asian genetic variation. These individuals may represent either unsampled wild populations from Central Asian transitional zones historically connecting eastern and western lineages, or captive-bred admixture between Asian and European genetic stocks. These findings underscore the need to extend the study to other regions of the Saker Falcon’s range; particularly northern/eastern Kazakhstan, areas east of the Caspian Sea, Western Kazakhstan, Uzbekistan, and Turkey. Such research is essential for developing effective conservation strategies and safeguarding the genetic diversity of wild populations.

### 2. Insights into the genetic status of captive saker falcons

Our analyses reveal striking contrasts between captive and wild saker falcons in both genetic diversity and inbreeding levels. Captive birds present both reduced observed heterozygosity (HOBS = 0.126 ± 0.008 vs. 0.143 ± 0.022 in wild conspecifics) and critically elevated inbreeding coefficients (FROH = 0.352 ± 0.024 vs. 0.131 ± 0.060) compared to their wild counterparts (Table 2; Figure 3). Such pronounced inbreeding is particularly concerning given established fitness consequences previously observed in birds. Flanagan et al. (2021) documented a 69% reduction in recruitment success in Hawaiian crow with pedigree-based inbreeding (F) above 0.098, while Duntsch et al. (2023) reported declines in juvenile survival of a New Zealand passerine at genome-derived FROH thresholds as low as 0.24. Notably, the mean FROH in our captive population (0.352) far exceeds both thresholds, suggesting severe risks to reproductive success and long-term viability, consistent with patterns of inbreeding depression observed in natural populations (Frankham et al., 2010). Together, these results demonstrate that inadequate genetic management of this captive population has accelerated genomic erosion, severely limiting its utility for conservation purposes.

Relatedness analyses reveal elevated and upwardly biased values (RAB) across the population (Figure 4). Specifically, 18.2% of captive dyads exceeded our threshold identifying related individuals (RAB > 0.154), indicating a functionally small breeding flock. The large spread with extreme values (RAB > 0.5) can reflect multi-generational sib-mating, consistent with the high inbreeding levels observed. Unlike standard relatedness metrics, RAB explicitly accounts for inbreeding bias (Korneliussen & Moltke, 2015), highlighting the genetic vulnerability observed in captive sakers. This vulnerability probably stems from three factors: (1) limited founders, (2) no recent genetic input, and (3) sub-optimal pairings. Due to high inbreeding and relatedness, these captive Sakers are of limited value for conservation unless supplemented through outcrossing with wild-origin individuals (Lacy, 1994; Rabier et al., 2020).

### 3. Recommendations for the genetic management of captive sakers

Our genomic assessment revealed that the potential founders showed hybridisation and severe inbreeding, highlighting the importance of systematic genetic screening prior to breeding and release, and revealing the critical challenges for conservation breeding. First, 19 birds (61%) showed evidence of introgression (three with gyrfalcons, 16 with an unknown species) and should be excluded from any conservation breeding programme. Of the remaining 15 pure sakers, 10 were genetically associated with Mongolia, while five may represent an unsampled Central Asian population. Alarmingly, all captive birds exhibited severe inbreeding (*FROH* > 0.3), with 18% of dyads being closely related (8.9% in Mongolian-associated birds, and three out of five birds in the subgroup 3). These findings necessitate a management approach consisting of: (1) removing introgressed individuals, (2) carefully pairing Mongolian-associated birds to limit further inbreeding, and (3) sourcing wild Central Asian individuals, both to validate the unsampled lineage and augment genetic diversity if confirmed. Without such interventions, this captive population would have limited, if any, interest for conservation purposes.

### 4. Conclusion

This study highlights the transformative potential of such a genomic framework in ex-situ conservation, as demonstrated through the comprehensive assessment of captive Saker Falcons. The implementation of standardised genomic protocols, encompassing introgression detection, population assignment, inbreeding levels and relatedness analysis, should become mandatory to ensure an efficient genetic management of conservation programme while guiding pairing decisions to maintain diversity and avoid inbreeding. While challenges such as cost and technical capacity persist, the integration of these tools addresses critical gaps in traditional management, preventing inadvertent selection of unfit individuals. Ultimately, this work establishes that genomic approaches are indispensable for evidence-based conservation, offering a replicable model for other threatened taxa facing similar genetic challenges.

## Acknowledgements

We are grateful to HH Sheikh Mohamed bin Zayed Al Nahyan, Crown Prince of Abu Dhabi and Founder of the IFHC, HH Sheikh Theyab Bin Mohamed Al Nahyan, Chairman of the IFHC, and HE Mohammed Ahmed Al Bowardi, Deputy Chairman, for their support. This study was conducted under the guidance of Reneco International Wildlife Consultants LTD, a company that manages the IFHC’s conservation programs, such as the National Avian Research Centre. Special thanks to Muhammad Sajid Saleem for creating the distribution map.

## Data availability statement

The data is accessible in the European Nucleotide Archive (ENA) at EMBL-EBI under accession number (PRJEB91371).

## Author contributions

T.B.H. and L.L. conceived and designed the study. A.B. and G.L. provided samples and associated metadata. X.V. performed laboratory work. T.B.H. conducted bioinformatic and population genomic analyses. T.B.H. led manuscript writing, with critical intellectual input from all co-authors. All authors reviewed, edited, and approved the final manuscript.

---

**Table S1.**
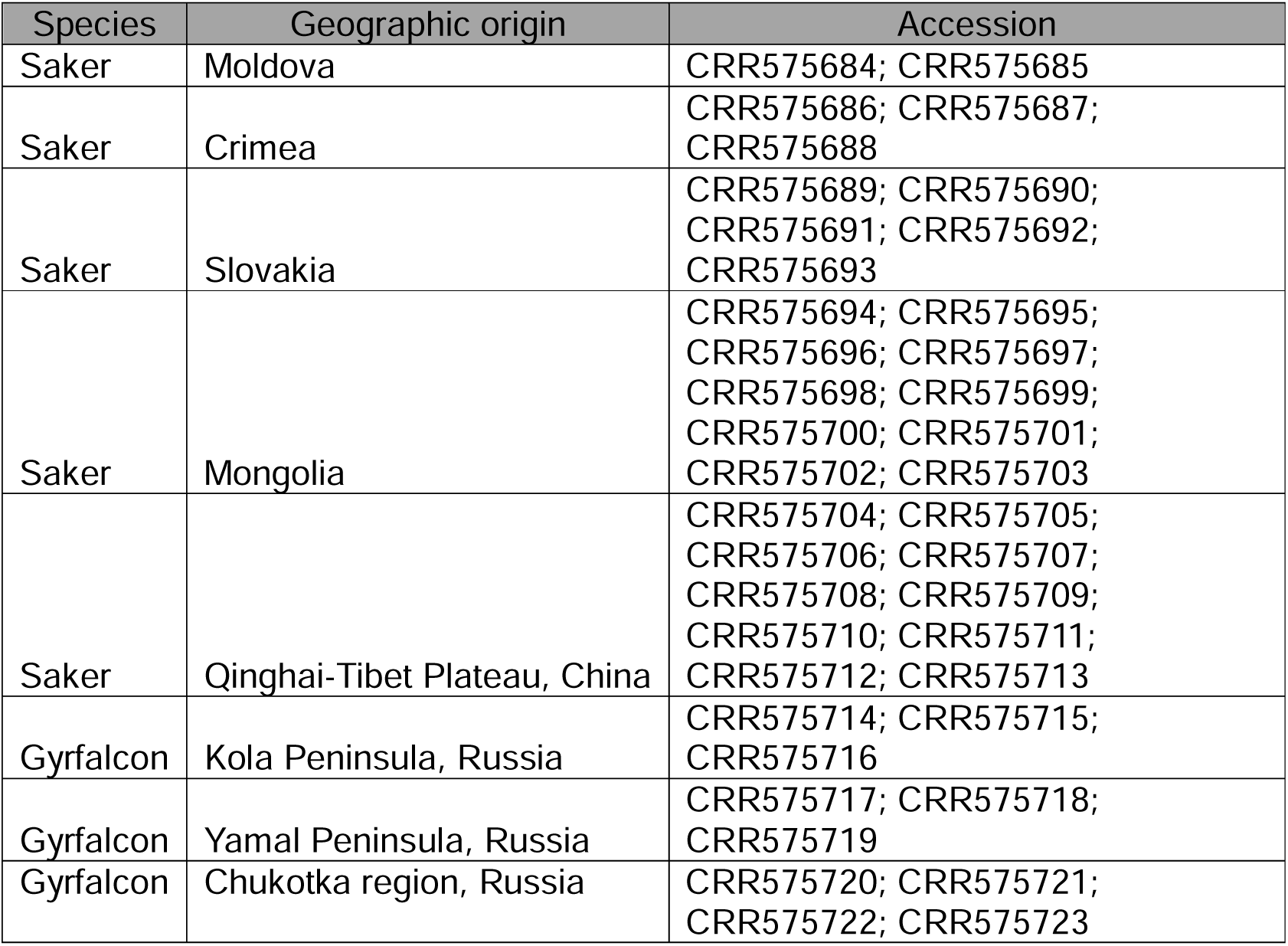
Sample details of the saker falcons (n = 30) and gyrfalcons (n = 10) used as reference populations (Fu et al. 2022).

| Species | Geographic origin | Accession |
| --- | --- | --- |
| Saker | Moldova | CRR575684; CRR575685 |
| Saker | Crimea | CRR575686; CRR575687;<br>CRR575688 |
| Saker | Slovakia | CRR575689; CRR575690;<br>CRR575691; CRR575692;<br>CRR575693 |
| Saker | Mongolia | CRR575694; CRR575695;<br>CRR575696; CRR575697;<br>CRR575698; CRR575699;<br>CRR575700; CRR575701;<br>CRR575702; CRR575703 |
| Saker | Qinghai-Tibet Plateau, China | CRR575704; CRR575705;<br>CRR575706; CRR575707;<br>CRR575708; CRR575709;<br>CRR575710; CRR575711;<br>CRR575712; CRR575713 |
| Gyrfalcon | Kola Peninsula, Russia | CRR575714; CRR575715;<br>CRR575716 |
| Gyrfalcon | Yamal Peninsula, Russia | CRR575717; CRR575718;<br>CRR575719 |
| Gyrfalcon | Chukotka region, Russia | CRR575720; CRR575721;<br>CRR575722; CRR575723 |

**Table S2.** Sequence quality of saker falcon samples from the SKHBC programme before and after trimming and filtering using FASTP v0.20.0. The average read depth is 23.7× across the 31 samples.

| *Individual ID* | *Raw reads* | *Size (GB)* | *Filtered reads* | *filtered reads (%)* | *Duplication rate* | *Insert size peak* | *Read depth* |
| --- | --- | --- | --- | --- | --- | --- | --- |
| F11K00001 | 293,639,774 | 20 GB | 282,122,392 | 96% | 0.09988933 | 151 | 27.3 |
| F12K00001 | 243,101,616 | 17 GB | 235,402,004 | 97% | 0.1006225 | 230 | 22.9 |
| F13K00002 | 261,703,670 | 18 GB | 254,933,184 | 97% | 0.12595239 | 271 | 24.4 |
| F14K00001 | 290,753,160 | 20 GB | 282,170,114 | 97% | 0.11211918 | 239 | 27.3 |
| F14K00002 | 230,447,374 | 16 GB | 223,042,754 | 97% | 0.09391843 | 151 | 21.4 |
| F15K00002 | 217,851,642 | 16 GB | 210,579,632 | 97% | 0.096838569 | 151 | 21.4 |
| F16K00001 | 294,450,216 | 22 GB | 286,394,818 | 97% | 0.03978355 | 261 | 30.3 |
| F16K00002 | 247,301,954 | 19 GB | 240,345,154 | 97% | 0.04140351 | 261 | 25.9 |
| F16K00003 | 257,877,244 | 19 GB | 250,901,864 | 97% | 0.04341457 | 261 | 25.9 |
| F17K00003 | 301,990,272 | 21 GB | 292,503,702 | 97% | 0.11826917 | 151 | 28.8 |
| F20K00002 | 236,835,718 | 18 GB | 230,124,274 | 97% | 0.03476558 | 261 | 24.4 |
| F20K00003 | 263,148,908 | 18 GB | 253,156,388 | 96% | 0.11197183 | 151 | 24.4 |
| F20K00004 | 340,428,064 | 26 GB | 332,010,338 | 98% | 0.04699982 | 261 | 36.2 |
| F20K00005 | 254,693,636 | 18 GB | 245,828,674 | 97% | 0.09580919 | 151 | 24.4 |
| F20K00006 | 245,077,920 | 17 GB | 236,590,508 | 97% | 0.09600089 | 151 | 22.9 |
| F20K00007 | 209,595,608 | 15 GB | 203,892,460 | 97% | 0.10682585 | 151 | 20.0 |
| F20K00008 | 280,344,816 | 21 GB | 271,755,438 | 97% | 0.04211539 | 261 | 28.8 |
| F11M00074 | 196,498,730 | 14 GB | 190,016,776 | 97% | 0.09475669 | 271 | 18.5 |
| F12M00107 | 221,464,292 | 16 GB | 213,579,098 | 96% | 0.10980367 | 261 | 21.4 |
| F12M00109 | 285,554,446 | 20 GB | 275,519,414 | 96% | 0.12188499 | 151 | 27.3 |
| F13M00018 | 206,612,838 | 15 GB | 200,901,696 | 97% | 0.10686098 | 271 | 20.0 |
| F13M00054 | 217,846,606 | 16 GB | 210,642,468 | 97% | 0.10050354 | 271 | 21.4 |
| F13M00082 | 193,239,332 | 14 GB | 184,299,410 | 95% | 0.09779283 | 271 | 18.5 |
| F13M00098 | 209,371,782 | 14 GB | 198,908,986 | 95% | 0.09052092 | 151 | 18.5 |
| F13M00103 | 296,340,866 | 20 GB | 283,785,174 | 96% | 0.11069308 | 196 | 27.3 |
| F13M00140 | 203,216,064 | 14 GB | 197,519,518 | 97% | 0.09448587 | 270 | 18.5 |
| F13M00149 | 190,450,836 | 14 GB | 184,007,718 | 97% | 0.08714687 | 237 | 18.5 |
| F14M00022 | 234,880,608 | 16 GB | 226,452,910 | 96% | 0.1031364 | 151 | 21.4 |
| F14M00063 | 230,662,912 | 16 GB | 223,019,528 | 97% | 0.11216896 | 262 | 21.4 |
| F14M00065 | 248,340,190 | 17 GB | 241,488,208 | 97% | 0.10060782 | 151 | 22.9 |
| F14M00077 | 231,993,478 | 16 GB | 224,306,852 | 97% | 0.09692249 | 151 | 21.4 |

**Figure S1.**
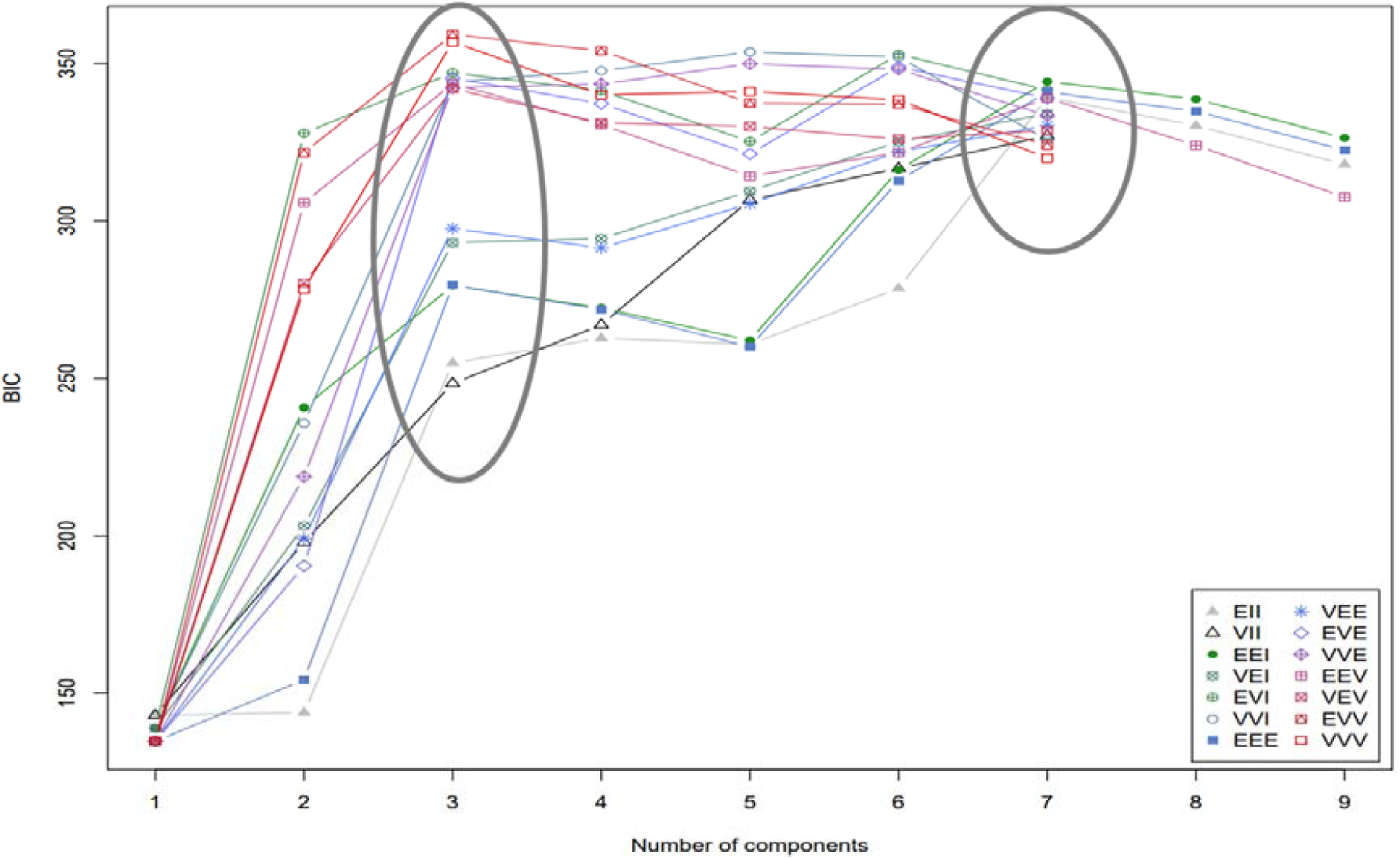
Model selection for genetic clustering using MCLUST. Bayesian Information Criterion (BIC) values for 14 covariance models across K = 1–9 clusters, calculated from principal component analysis (PCA) coordinates. The optimal solution (EVV model, K = 3; circled) showed the highest BIC, with K = 7 also demonstrating biologically meaningful structure. Ten models failed to converge at K ≥ 8.

